# The YcnI protein from *Bacillus subtilis* contains a copper-binding domain

**DOI:** 10.1101/2021.06.11.448128

**Authors:** Madhura S. Damle, Stephen C. Peters, Veronika A. Szalai, Oriana S. Fisher

**Author notes:** Corresponding author: Oriana S. Fisher.

## Abstract

Bacteria require a precise balance of copper ions to ensure that essential cuproproteins are fully metalated while also avoiding copper-induced toxicity. The Gram positive bacterium *Bacillus subtilis* maintains appropriate copper homeostasis in part through its *ycn* operon. The *ycn* operon comprises genes encoding three proteins: the putative copper importer YcnJ, the copper-dependent transcriptional repressor YcnK, and the uncharacterized DUF1775 domain-containing YcnI. DUF1775 domains are found across bacterial phylogeny and bioinformatics analyses indicate that they frequently neighbor domains implicated in copper homeostasis and transport. Here, we investigated whether YcnI can interact with copper and, using electron paramagnetic resonance (EPR) and inductively-coupled plasma-mass spectrometry (ICP-MS) find that it can bind a single Cu(II) ion. We determine the structure of both the apo and copper-bound forms of the protein by X-ray crystallography, uncovering a copper binding site featuring a unique mono-histidine brace ligand set that is highly conserved among DUF1775 domains. These data suggest a possible role for YcnI as a copper chaperone and that DUF1775 domains in other bacterial species may also function in copper homeostasis.

## Introduction

Copper is an essential cofactor for many enzymes, but in high quantities it can also have deleterious effects on cellular viability due to formation of reactive oxygen species. In nearly all bacterial species, a suite of proteins maintains an appropriate balance of copper by regulating its homeostasis and mediating its transport in and out of the cell (1, 2). Copper efflux is usually carried out by Cu(I) exporting P-type ATPases, which often are further assisted in this process by additional proteins. For example, in the Gram positive *Bacillus (B*.*) subtilis*, the CopA Cu-dependent ATPase exports copper with the aid of the CopZ chaperone (3–5). The *copZA* operon is further regulated by CsoR, a copper(I) sensing repressor, that binds to the operator region in the absence of copper (6, 7).

Bacterial copper acquisition, on the other hand, has remained more elusive. Data from a number of different organisms, however, have converged to suggest that the CopD domain functions as a membrane-bound copper importer. Proteins containing CopD domains have been found to play an essential role in conferring copper resistance in a wide range of microorganisms including *B. subtilis* (8, 9), *Pseudomonas syringae* (10, 11), *Acinetobacter baumannii* (12), and *Bradyrhizobium japonicum* (13). These and other microbiological and transcriptomic studies all strongly point towards a role for this domain in copper uptake (8, 9, 12, 14–16). Many of proteins with CopD domains, however, are encoded by operons that include additional genes such as *copC* and *pCu*_*A*_*C* that whose protein products have also been implicated in copper homeostasis (17, 18), as well as other yet to be characterized proteins of unknown function.

One notable example is the *ycnKJI* operon in *B. subtilis* that encodes the YcnK, YcnJ, and YcnI proteins. YcnK is a Cu-dependent transcriptional repressor that uses a helix-turn-helix (HTH) domain to bind to an intergenic region upstream of the *ycn* operon (9), controlling its expression. Because loss of *ycnJ* results in a reduction of intracellular copper levels (8, 9), the YcnJ protein has been proposed to serve as a copper importer. As further evidence for such a role, YcnJ is a fusion protein comprised of an extracellular CopC domain of the C_0-1_ type that typically binds Cu(II), a membrane-bound CopD domain, and a C-terminal YtkA domain of unknown function (18).

The third protein encoded by the *ycn* operon, YcnI, remains much more poorly understood. It includes a <u>D</u>omain of <u>U</u>nknown <u>F</u>unction 1775 (DUF1775) and a C-terminal transmembrane (TM) helix (19, 20). Its biological function has not been determined, but its frequent association with CopC and CopD suggests that it likely is also involved in copper homeostasis (9, 13, 17, 21, 22). Furthermore, YcnI is homologous to the PcuD protein encoded by the *pcuABCDE* operon from *Bradyrhizobium japonicum* that also appears to play a role in copper acquisition (13). Studies of the *ycn* operon have alluded to a potential role for YcnI in copper homeostasis (9). The lack of biochemical information about this protein and the absence of a recognizable metal-binding motif in its amino acid sequence, however, have impeded further advances towards elucidating its functional role.

Here, we investigate a link between YcnI and copper. We find that YcnI homologs are present in a wide range of bacterial species, with particularly high representation within the firmicutes, actinobacteria, and proteobacteria phyla, and that genes encoding DUF1775 domains frequently occur in close proximity to other copper-related genes. We recombinantly produce the soluble domain of *Bacillus subtilis* YcnI (*Bs-*YcnI) *in vitro* and demonstrate that it binds Cu(II) in a 1:1 stoichiometry. To investigate the structure of the protein and to further probe the coordination geometry of the copper site, we determine crystal structures of the protein in both the apo and Cu-bound states. The latter reveals an unusual copper binding motif that we term the mono-histidine brace due to its similarities to the canonical histidine brace coordination site used by some monooxygenases and other copper binding proteins (17, 23, 24). The metal binding ligands identified in the crystal structure are strictly conserved in the majority of YcnI sequences, suggesting a role for YcnI homologs in copper homeostasis and trafficking in many bacterial species.

## Results

### Genomic analyses suggest an association between DUF1775 domains and copper-binding proteins

To investigate the role of the DUF1775 domain in bacteria, we conducted a large-scale bioinformatics analysis of DUF1775 domains. We mined the Joint Genome Institute-Integrated Microbial Genomes (JGI-IMG) database (25) for sequences that match the associated protein family (PFAM), pfam07987, identifying over 10,000 individual sequences that we used to construct a sequence similarity network. Notably, DUF1775 domains appear throughout bacterial phylogeny, occurring in 9354 unique genomes and within both Gram positive and Gram negative organisms (**Fig. S1A**). The domain is most highly represented in the proteobacteria, actinobacteria, and firmicutes phyla, as well as a significantly smaller representation in deinococci. The majority (≈ 90 %) of these sequences contain a single DUF1775 domain, while a subset of the sequences from proteobacterial species fuse the DUF1775 domain to a <u>p</u>eriplasmic <u>Cu_A_ c</u>haperone (PCuAC) domain (**Fig. S1B**). This corroborates previous studies of *DUF1775* genes that noted a co-occurrence with *pCuAC* and other genes such as *copC* and *copD* that encode copper-related proteins (9, 13, 17, 22).

To understand more about how the *B. subtilis* YcnI protein compares to other proteins with DUF1775 domains, we narrowed in on the cluster of DUF1775 sequences in which *B. subtilis* YcnI is found. *Bs*YcnI appears most closely related to DUF1775 domains found in other members of the *Bacillus* genus, as well as other firmicutes, most notably *Listeria* and *Paenibacillus* (**Fig. 1A**). Of these proteins, ≈ 40 % (270 out of a total 606 sequences) neighbor a CopCD-encoding gene, including those like *ycnJ* (**Fig 1B**). Approximately 20 % of these genes also are near a YcnK-like or HTH-containing protein, comprising what, in *B. subtilis*, has been termed the *ycn* operon (**Fig. 1C**). Despite its frequent proximity to copper-associated domains, however, YcnI does not contain any known conserved metal binding motifs. Examination of the sequence logo for the DUF1775 family reveals one very highly conserved histidine residue (present in 10393 of 10647 sequences) near the N-terminus, but no other obvious conserved residues or motifs for metal binding.

**Figure 1.**
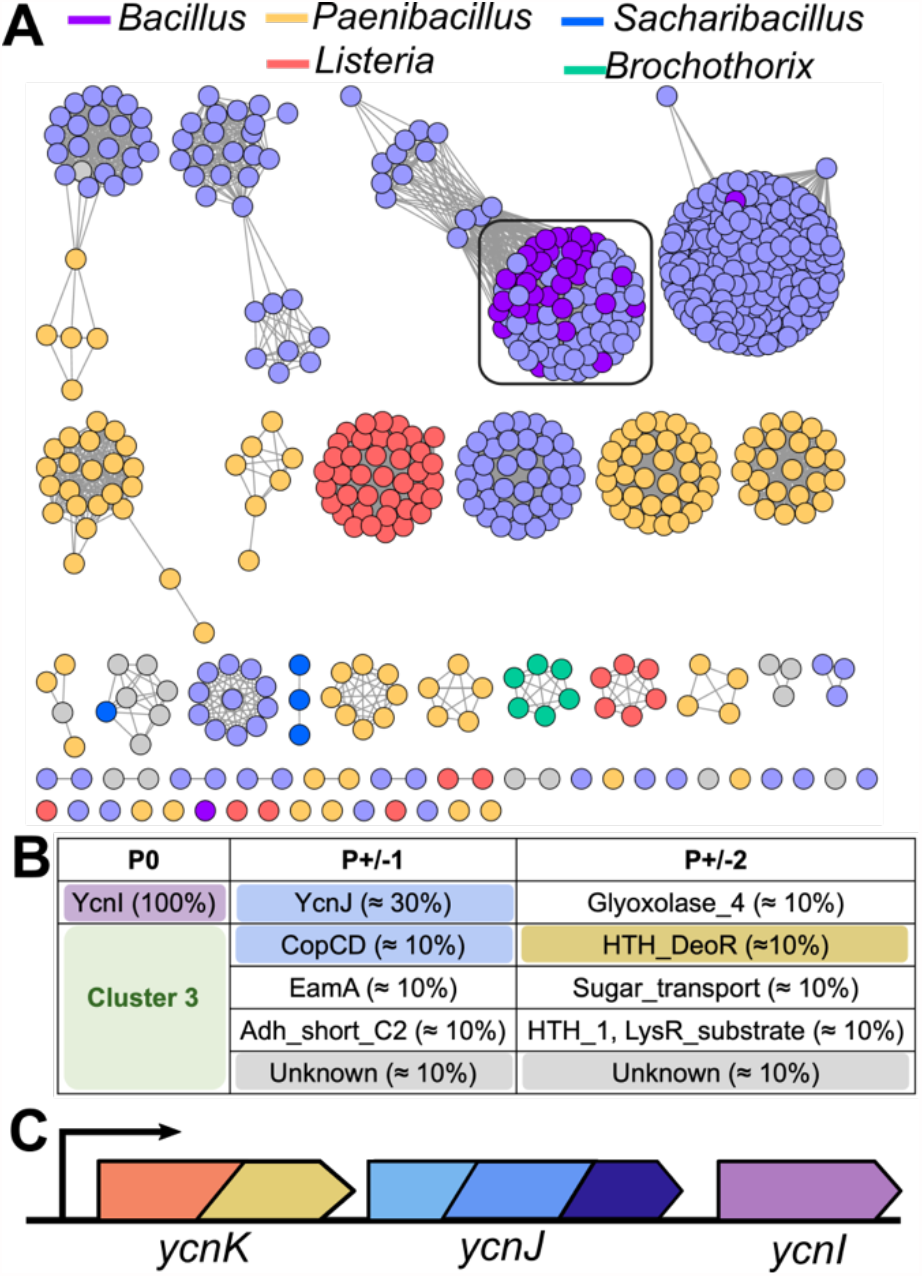
Bioinformatics analysis of DUF1775 sequences most closely related to *B. subtilis* YcnI. **A)** Sequence similarity network for DUF1775 sequences in the same cluster as YcnI colored by genus, alignment score cut-off of 70. Sequences are colored by taxonomy, and the cluster containing the *B. subtilis* YcnI is boxed. **B)** Genome neighbors within two positions of the sequences shown in A. **C)** Architecture of the *ycn* operon.

### BsYcnIΔC binds Cu(II)

Because our bioinformatics data suggested a link between YcnI and proteins such as CopC, CopD, and PCu_A_C involved in copper homeostasis, regulation, and trafficking, we next sought to determine if YcnI binds copper. The *B. subtilis* YcnI protein contains a predicted signal sequence N-terminal to the DUF1775 domain, suggesting that endogenously, the DUF1775 domain is localized extracellularly and is tethered to the membrane by the C-terminal TM helix (**Fig. S1B**). Secondary structure predictions and sequence alignments suggested that upon cleavage of the signal peptide, the N-terminal residue would be the well-conserved histidine. We hypothesized that this histidine might serve a role as a metal ligand, analogous to how an N-terminal histidine residue coordinates copper ions in other bacterial proteins including lytic polysaccharide monooxygenases (LPMOs) (23, 26–31), CopC (18, 24, 32–34), the periplasmic copper-A chaperone (PCu_A_C) domains of PmoF (17), and the Cu_B_ site of particulate methane monooxygenase (pMMO) (35).

To generate the YcnI protein with the native N-terminal histidine residue intact, we engineered a small ubiquitin-like modifier (SUMO)-tagged construct of the DUF1775 domain of YcnI (*Bs*YcnIΔC). We expressed this construct in *E. coli* and purified it to homogeneity after cleavage of the SUMO tag. To investigate the possibility that *Bs*YcnIΔC could bind copper, we incubated the purified protein with different stoichiometric ratios of CuSO_4_ and measured the concentration of bound copper by inductively coupled plasma-mass spectrometry (ICP-MS) after removing unbound metal using a desalting column. We find that addition of either 1 equivalent or 2 equivalents resulted in 0.98 ± 0.09 Cu ions bound per protein, indicating that the protein binds the metal ion in a 1:1 stoichiometry when either 1 or 2 equivalents of the metal are added. The as-isolated apo samples did not have significant copper content (0.081 ± 0.06 Cu ions per protein monomer) (**Table 1**).

**Table 1.** YcnIΔC copper binding stoichiometry.

| **Sample** | **Mole equivalent Cu: protein monomer** |
| --- | --- |
| *Bs*-YcnIΔC, apo | $0.08 \pm 0.06$ |
| *Bs*-YcnIΔC, Cu-loaded* | $0.98 \pm 0.09$ |

\*Samples were incubated in a one to one ratio with CuSO<sub>4</sub> prior to desalting. Uncertainties are standard deviations, calculated from 3 (apo) and 5 (Cu-loaded) independent replicates.

To further investigate the copper coordination environment and to determine the oxidation state of the metal, we performed electron paramagnetic resonance (EPR) spectroscopy on the Cu-loaded protein. The axial EPR spectrum clearly shows characteristic hyperfine splitting pattern in the g_||_ region indicative of Cu(II) (g_||_ ≈ 2.26 and A_||_ ≈ (17.3 ±0.4) mT; g_⊥_ ≈ 2.06; see Materials & Methods section for uncertainty analysis) (**Fig. 2**). The g_||_ and A_||_ values suggest that the Cu(II) coordination environment includes a total of four ligands comprised of either 4 nitrogen or 2 nitrogen and 2 oxygen (36). Consistent with the ICP-MS data, the apo protein exhibited a weak Cu(II) EPR signal indicating the presence of a small amount of bound copper. Together, these data show that *Bs*YcnIΔC binds Cu(II) in a 1:1 stoichiometry.

**Figure 2.**
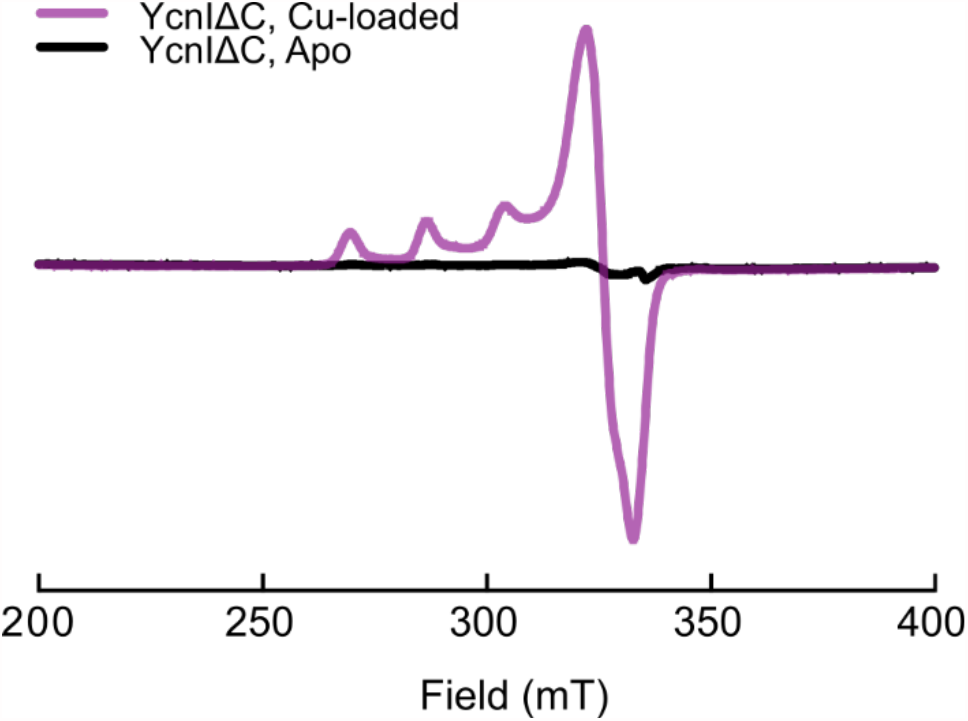
*Bs-*YcnI binds Cu(II). EPR spectra of Cu-bound *Bs*YcnIΔC (purple) and apo *Bs*YcnIΔC (black). nstrument settings: temperature 10.0 K ± 0.1 K; microwave power 0.47 mW; modulation amplitude, 0.5 mT; modulation frequency, 100 kHz; conversion time, 88 ms; 2048 points, 16 scans.

### BsYcnIΔC adopts a cupredoxin fold and binds copper at an N-terminal site

We next wanted to determine precisely where the Cu(II) binding site is in the protein as well as whether interactions with the metal induce any conformational changes, so we initiated structural studies of *Bs*YcnIΔC. We crystallized both the apo and copper-bound forms of the protein and determined their structures to 0.20 nm (2.05 Å) and 0.21 nm (2.11 Å) resolution, respectively (**Fig. 3, Table 2**). Overall, *Bs*YcnIΔC adapts a cupredoxin fold featuring the characteristic Greek key β barrel shared among many bacterial copper-binding proteins (37).

**Table 2.** Crystallographic data collection and refinement statistics.

| **Crystal** | **Cu *Bs-YcnI*ΔC** | **Apo *Bs-YcnI*ΔC** |
| --- | --- | --- |
| **PDB accession code** | **7MEK** | **7ME6** |
| **Data Collection** |  |  |
| Wavelength (Å)<sup>†</sup> | 1.377 | 0.979 |
| Space Group | *P*6<sub>3</sub>22 | *P*6<sub>3</sub>22 |
| Cell Dimensions |  |  |
| a, b, c (Å) | 90.2, 90.2, 209.1 | 90.4, 90.4, 208.4 |
| α, β, γ (°) | 90, 90, 120 | 90, 90, 120 |
| Resolution (Å) | 30-2.11 (2.16-2.11)* | 39.1-2.05 (2.11-2.05)* |
| *R*<sub>meas</sub> (%) | 13.8 (152.7)* | 13.8 (134.5)* |
| *I* / *σI* | 9.64 (2.64)* | 12.06 (1.77)* |
| Completeness (%) | 99.6 (98.0)* | 99.8 (97.4)* |
| Redundancy | 5.3 (5.3)* | 11.1 (8.1)* |
| **Refinement** |  |  |
| Resolution (Å) | 29.52-2.11 (2.14-2.11)* | 38.48-2.05 (2.10-2.05)* |
| No. of reflections | 54321 | 32328 |
| *R*<sub>work</sub>/*R*<sub>free</sub> (%) | 19.5 / 21.7 (30.8 / 30.9)* | 19.6 / 21.9 (26.3 / 29.3)* |
| Residue range built | A: 27-155, B: 27-153 | A: 27-155, B: 27-154 |
| No. of atoms |  |  |
| Protein | 2027 | 2040 |
| Ligand/ion | 2 Cu | 2 malonate |
| Water | 125 | 252 |
| **Model quality** |  |  |
| *B*-factors (Å<sup>2</sup>) |  |  |
| Overall | 47.05 | 48.16 |
| Protein | 46.84 | 47.72 |
| Ligand/ion | 54.78 | 71.61 |
| Water | 50.35 | 50.36 |
| r.m.s.d., bond lengths (Å) | 0.002 | 0.007 |
| r.m.s.d., bond angles (°) | 0.526 | 0.98 |
| Ramachandran favored/<br>allowed/outliers (%) | 97.33/2.78/0 | 98.81/1.19/0 |

<sup>†</sup>1 Å = 0.1 nm \*Values shown in parentheses are for the highest resolution shell.

**Figure 3.**
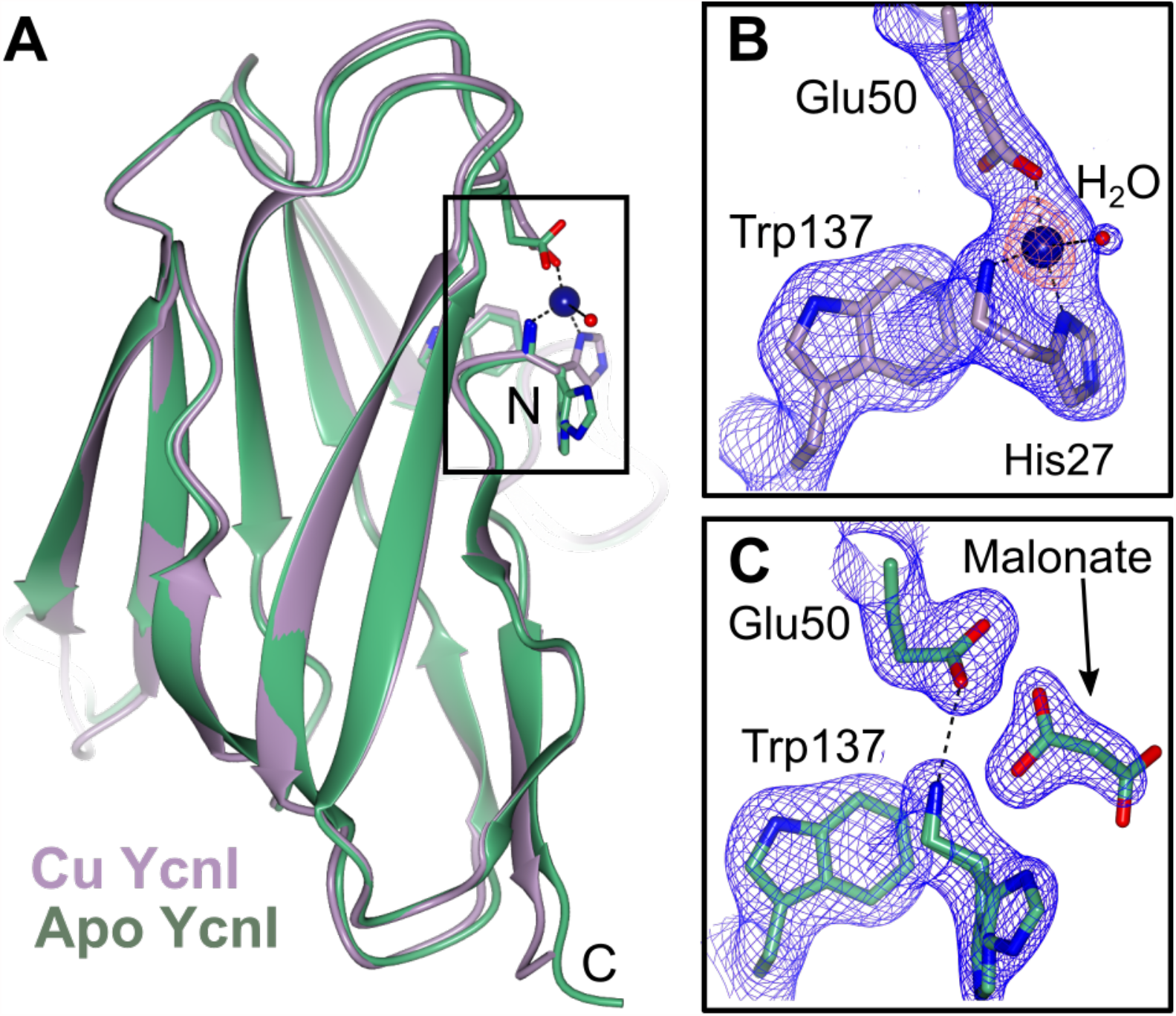
Crystal Structures of Cu-bound and apo *Bs*-YcnIΔC. **A)** Superposition of the Cu-bound *Bs*YcnIΔC (purple) and the apo *Bs*YcnIΔC (green). **B)** Close-up view of boxed region in the Cu-bound structure. The Cu ion is shown as a blue sphere. The 2*F*_o_*-F*_c_ map is shown in blue mesh (contoured to 1.5α) and the anomalous density map in orange mesh (contoured to 5α). Bonds are indicated by dashed lines. **C)** Close-up view of boxed region in the apo structure, including the malonate ion in grey. The 2*F*_o_*-F*_c_ map is shown in blue mesh (contoured to 1.5α).

In the structure of the copper-bound structure, we observed a strong peak in the anomalous difference map that we modeled as a single Cu ion near the N-terminus of the protein (**Fig. 3B**). The metal ion is coordinated both by the amino terminal nitrogen and the δ-nitrogen of His27 at a distance of 0.20 nm (2.0 Å), as well as the ε_2_-oxygen of Glu50 at a distance of 0.21 nm (2.1 Å). Both the ε_1_-oxygen of Glu50 and a water molecule are more weakly coordinating from distances of 0.30 nm (3.0 Å) and 0.35 nm (3.5 Å) away, respectively (**Fig. 3B**). These ligands correlate well with the 2N2O coordination suggested by the EPR data, with the two nitrogen ligands deriving from His27 and the two oxygen ligands from Glu50 and a water. In addition, one of the highly conserved tryptophan residues (Trp137) further cushions the metal ion by engaging in cation-π interactions from a distance of 0.34 nm (3.4 Å) away. The fact that the N-terminal histidine is highly conserved across DUF1775 domains (**Fig. S2)** further supports the idea that it plays a vital role in binding to copper.

The fold of the apo structure of *Bs*YcnIΔC is very similar to that of the Cu-bound protein with a root mean square deviation (RMSD) of 0.059 nm (0.59 Å) over 127 C_α_ (**Fig. 3A**). The primary differences are in the N-terminal region near the Cu-binding site. Instead of the His27 and Glu50 side chains turning in toward one another as they do in the Cu-bound form, in the apo form, these residues are oriented away from one another with the N-terminus directly hydrogen bonding to the side chain of Glu50 (**Fig. 3B**,**C**). In one of the two chains of the asymmetric unit, we observe two alternate conformations for His27, suggesting that this side chain displays a degree of flexibility. In our apo structure, we also observe a malonate molecule from the cryoprotectant solution near the N-terminus (**Fig. 3C**), indicating that this region of the protein is solvent accessible.

### Bs YcnI employs a mono-histidine brace motif to bind copper

The *Bs*YcnIΔC crystal structure reveals a unique metal binding site, with features reminiscent of two previously characterized Cu-binding sites (**Fig. 4A**). Most strikingly, the copper coordination observed in the crystal structure of *Bs*YcnIΔC most closely resembles the histidine brace motif. The His-brace motif was initially identified in LPMOs, copper-binding enzymes that oxidatively degrade cellulose and chitin (38). In LPMOs, the His-brace motif often serves a catalytic role. Although there are differences among the active sites, LPMOs typically use a 3N T-shaped configuration and occasionally feature a phenylalanine near the active site (38) **(Fig. 4B)**. Recently, an LPMO protein from *Laetisaria arvalis* (LaX325) was found to employ a His-brace for copper binding as well (**Fig. 4C**), despite an absence of catalytic activity (39). Beyond LPMOs, other non-enzymatic copper binding proteins including many members of the CopC family (**Fig. 4D**) of proteins involved in copper homeostasis (22, 34) and the PCu_A_C domains of the PmoF proteins encoded by some species of methanotrophic bacteria (40) (**Fig. 4E**) also use histidine braces to coordinate Cu ions. The histidine brace motifs in all of these proteins use an N-terminal histidine residue to provide two coordinating ligands to the metal: the amino terminus of the protein and one of the imidazole nitrogens of the histidine side chain (23). A second histidine residue provides a third nitrogen ligand, while often one or more oxygen ligands are further contributed by glutamate, aspartate, or water molecules.

**Figure 4.**
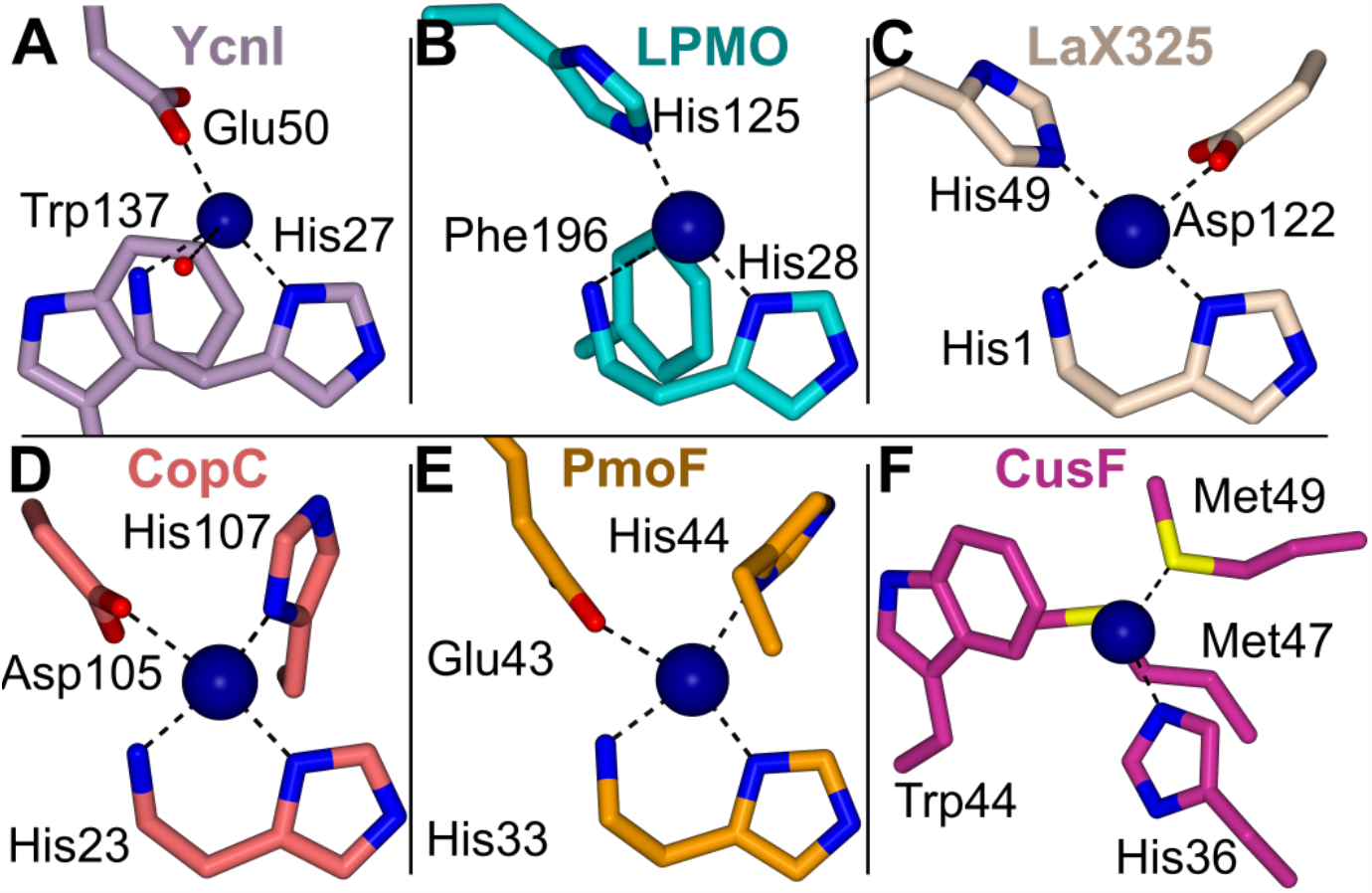
Comparison of *Bs*YcnI with other Cu-binding proteins. The Cu binding sites of **A)** *Bacillus subtilis* YcnI (this study) **B)** *Bacillus amyloliquefaciens* CBM33 LPMO (PDB ID: 2YOX), **C)** the LaX325 enzyme (PDB ID: 6IBH, chain A), **D)** *Methylosinus trichosporium* OB3b CopC (PDB ID: 5ICU), **E)** *Methylocystis sp*. Rockwell PmoF1 (PDB ID: 6P16), and **F)** *E. coli* CusF (PDB ID: 2VB2) shown as sticks with bonds indicated by dashed lines and coordinating residues highlighted.

Despite the marked similarities between the *Bs*YcnI Cu site and those of structurally characterized proteins that use a canonical His-brace motif, the second coordinating histidine residue is notably absent in *Bs*YcnI (**Fig. 4A**). The copper coordination site in *Bs*YcnI differs from these other proteins in that a glutamate residue completes the coordination sphere in lieu of a second histidine. Although the use of an N-terminal histidine residue as a metal ligand is a common feature among His-brace proteins, to our knowledge, all other such sites include the second coordinating histidine to form the T-shaped geometry. Our discovery of the coordination site in *Bs*YcnI introduces a new subcategory within histidine brace motifs. We term this motif the mono-histidine brace to differentiate it from the bis-histidine brace observed in other copper-binding proteins and enzymes.

The second unique feature of the *Bs*YcnI copper binding site is an adjacent tryptophan residue, Trp137, that is exceptionally well-conserved across YcnI homologs. Although tryptophan residues do not traditionally engage in interactions with copper ions, there are a few notable exceptions. The CusF copper chaperone uses a somewhat similar ligand geometry (**Fig. 4F)** (41) and in the MopE protein an oxidized tryptophan residue serves as a metal ligand (42). A recent study of CopG, a copper dependent oxidoreductase, also features a strictly conserved tryptophan residue in close proximity to one of the metal ions (43). The orientation of the tryptophan in YcnI is most similar to the that of the tryptophan in the Cu-binding site of CusF. Unlike the hydrophobic environment of the CusF tryptophan, however, Trp137 of *Bs*YcnI is located in a solvent-exposed region of the protein instead (44). Overall, the metal coordination site of *Bs-*YcnI appears to meld features of both the His-brace and the CusF-style copper centers using a mono-histidine brace motif.

### Copper binding residues are highly conserved in YcnI sequences

Although *Bs*YcnI and *B. japonicum* PcuD have been implicated in copper homeostasis (13, 20), a specific role for copper binding for other YcnI family members has not been investigated directly. To gain insights into whether other homologs could engage in copper interactions in a similar fashion to *Bs*YcnI, we revisited our bioinformatics analyses to determine whether the residues we identified as the mono-histidine brace motif are conserved among other DUF1775 family members as well. We aligned all sequences to the DUF1775 HMM and found that the His, Glu, and Trp residues are conserved in most of these proteins (≈ 60 %) (**Fig. 5, Fig. S2**). Of the remaining sequences, which predominantly are found in proteobacterial genomes, the majority have the His and Trp conserved but not the Glu, suggesting that these proteins may either have a distinct function or use a different ligand set. Overall, ≈ 95 % of the YcnI sequences have the histidine and tryptophan ligands fully conserved. These data strongly suggest that YcnI is a new player in bacterial copper homeostasis, beyond the *Bacillus subtilis* homolog investigated here.

**Figure 5.**
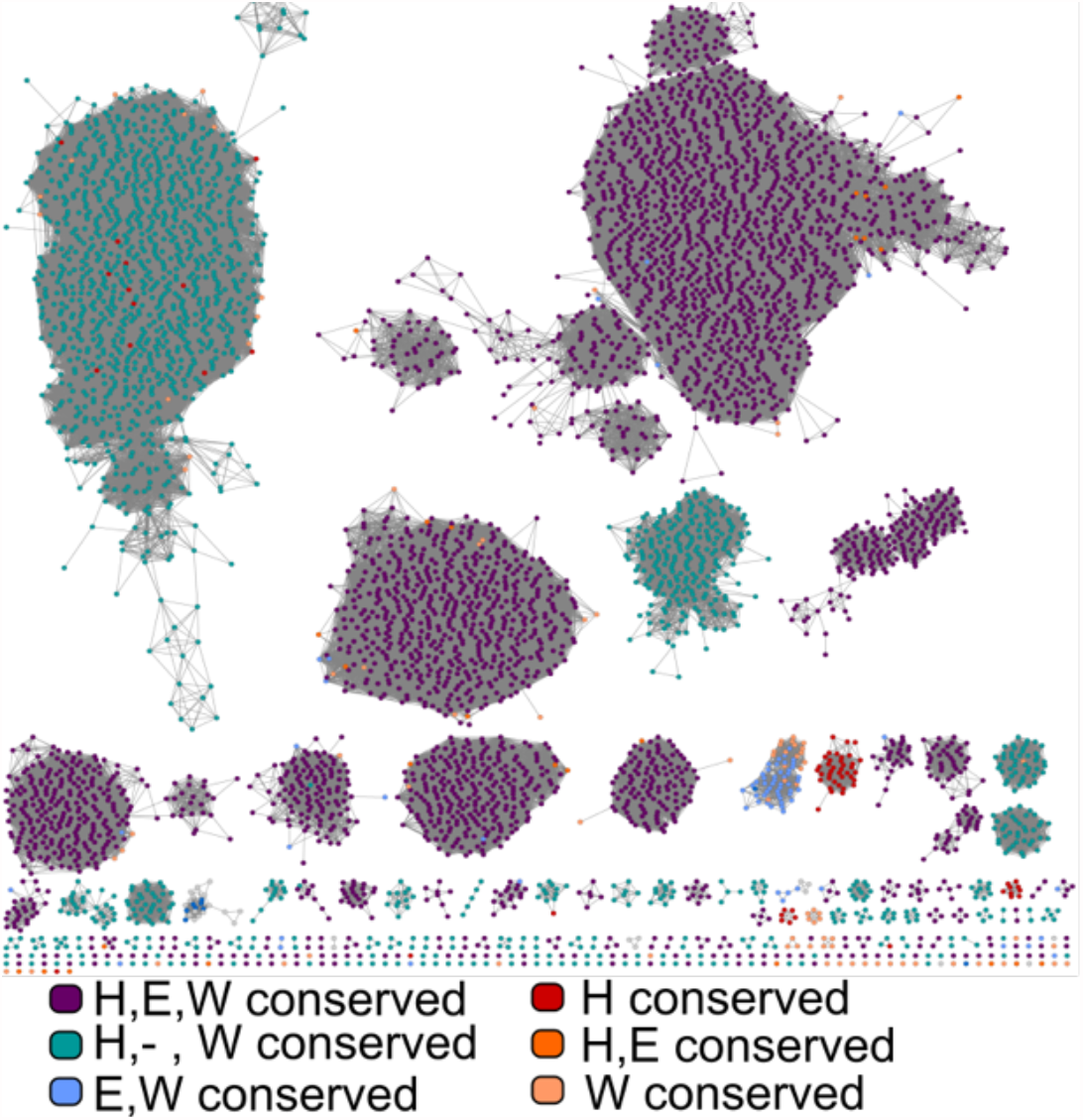
Sequence conservation in the DUF1775 family. Sequence similarity network for the DUF1775 family, colored by conservation of the Cu-binding ligands (His27, Glu50, Trp137) identified in *Bs*YcnI as generated through sequence alignment in ClustalO. Sequences with all three residues conserved are indicated as purple circles, those with only the His and Trp conserved in teal, those with only the Glu and Trp conserved in light blue, those with the His and Glu conserved in dark orange, those with only the Trp conserved in light orange, and those with only the His conserved in red.

## Discussion

The DUF1775 domain has been suggested to play a role in copper homeostasis or trafficking, and the *Bs-*YcnI protein specifically has been proposed to serve in such a capacity. Corroborating this idea, here we find that the distribution of DUF1775 domains frequently occurs among a significant number of bacterial species. Our spectroscopic data on the *B. subtilis* YcnI show that this domain binds a single Cu(II) ion, which crystallographic data further reveal to be coordinated by a solvent-accessible, N-terminal metal ion binding site. The specific ligands that coordinate the copper ion are highly conserved among a majority of members of the DUF1775 family. Together, these data strongly suggest that YcnI, and more broadly the DUF1775 domain, represent a new class of bacterial copper binding proteins.

*Bs*YcnIΔC binds to Cu(II) through the backbone nitrogen at the N-terminus, a nitrogen from the N-terminal histidine, an oxygen of a glutamate, and a water molecule. This mono-histidine brace coordination motif and the presence of a nearby tryptophan residue represent a distinct variation on the canonical bis-histidine brace motif, raising a number of possibilities for its biological role. LPMO enzymes that use a mononuclear copper site coordinated by a bis-histidine brace motif frequently exhibit oxidative properties. In these enzymes, the bis-histidine brace motif has been proposed to assist in the formation of stable, high-valent intermediates through deprotonation of the amino group (30). Studies investigating the copper-dependent monooxygenases have established a possible link between catalytic function and the bis-histidine brace (30, 45), although the CopC and PmoF proteins that also use the canonical histidine brace do not appear to have enzymatic activity (17, 22). The mono-histidine brace motif we have identified in *Bs*YcnIΔC, with its relatively low coordination number and a more solvent exposed copper site raise the possibility *Bs*YcnI may not use copper for catalysis and could potentially facilitate transfer of copper ions between proteins.

Our discovery of the mono-histidine brace copper coordination site represents an important step towards understanding *Bs*YcnI, but the specific biological function of this protein remains to be fully elucidated. Since Δ*ycnJ* demonstrates a growth defective phenotype in the absence of copper and is predicted to have a membrane bound domain, it has been suggested to play a role in copper homeostasis (20). One possibility is that *Bs*YcnI could act as a copper chaperone to the putative copper importing protein YcnJ. It may form a complex of high coordination number with the extracellular CopC domain of YcnJ which in turn could facilitate copper uptake via the CopD domain, a putative copper importer (9). Alternately, the absence of the second histidine ligand could result in reduced metal affinity for the mono-histidine brace site compared to the bidentate site in the CopC domain, perhaps promoting unidirectional transfer of copper ions to the importer.

Our bioinformatics analyses indicate that YcnI homologs are found in a wide variety of bacterial species, including a number of pathogenic strains like *Mycobacterium tuberculosis, Listeria monocytogenes, Bacillus anthracis*, and others. Many of these and other homologs also share the copper binding residues we identified in *Bs*YcnI. The existence of a subset of proteobacterial DUF1775 domains that are fused to PCu_A_C domains is also particularly intriguing in light of our findings that YcnI binds copper. It is possible that in such fusion proteins, the DUF1775 and PCu_A_C domains work in concert to maintain copper homeostasis. Because modulation of intracellular copper levels in *M. tuberculosis*, for example, can be exploited by the host immune system to combat invading pathogens (2, 46), it will be of particular importance to investigate a possible role for YcnI in bacterial copper resistance mechanisms.

## Materials and Methods

Certain commercial equipment, instruments, or materials are identified in this paper in order to specify the experimental procedure adequately. Such identification is not intended to imply recommendation or endorsement by the National Institute of Standards and Technology, nor is it intended to imply that the materials or equipment identified are necessarily the best available for the purpose.

### Bioinformatics analyses

To identify protein sequences that include DUF1775 domains, the IMG-JGI database was interrogated for genes encoding its corresponding PFAM (pfam07987) in finished bacterial genomes, resulting in the identification of 10646 sequences. To generate a sequence similarity network, these sequences were submitted to the EFI-EST server using Option C. Sequences with fewer than 100 amino acids were excluded from the analysis, and an alignment score of 50 and E-value cut-off of 5 were used to generate the network. Due to the large size, for all subsequent analysis the 100 % network (in which identical sequences are represented as a single node) was used. Metadata for each sequence including pfam identifiers and taxonomic information were extracted from IMG-JGI and imported into the network table in Cytoscape. A fasta file containing all sequences in the network was aligned to the hidden Markov model (HMM) for PFAM07987 using ClustalO to determine the conservation of the metal binding ligands. The identity of the amino acids at each of the positions corresponding to the metal binding ligands were added to the metadata table (**File S1**). To identify gene neighbors of DUF1775 sequences above, all genes within one position from each gene identified above were extracted from IMG-JGI (25). These data were then used to calculate the frequency at which different domains are found in the neighboring genes to the YcnI sequence. A fasta file for the sequences in the cluster containing YcnI was generated, aligned to the HMM using ClustalO (47, 48), and the resulting file was used to generate a sequence logo in SkyLign (49). All software used for bioinformatics analyses are open source.

### Construct design

DNA for the soluble domain (residues 27-155) of *Bacillus subtilis ycnI* (Uniprot ID: P94431) was synthesized into the pET28a+TEV vector using the NdeI and BamHI restriction sites. To generate the authentic N-terminal histidine residue, a His-SUMO tag was inserted immediately upstream. Briefly, the pET 28a+TEV vector was linearized using Primer Set 1 and the His-SUMO sequence was amplified using Primer Set 2 to generate matching overhangs (**Table S1**). The resulting DNA was assembled using Gibson Assembly to generate the HisSUMO-YcnIΔC plasmid.

### Protein expression and purification

The HisSUMO-YcnIΔC construct was transformed into BL21(DE3) cells. Overnight cultures were inoculated into Luria-Bertani media, and protein expression was induced by the addition of 1 mmol/L isopropyl β-d-1-thiogalactopyranoside at *A*_600_ ≈ 0.6. The cultures were then grown overnight at 20 °C and harvested by centrifugation at 737.5 rad/s (6000 x *g*) for 20 min. The pellet was resuspended in lysis buffer (150 mmol/L NaCl, 20 mmol/L 4-(2-hydroxyethyl)-1-piperazineethanesulfonic acid (HEPES), 20 mmol/L imidazole, pH 7.5) supplemented with 1 mmol/L DTT and 0.5 mmol/L PMSF prior to sonication. After sonication, the sample was centrifuged for 45 min at 1257.8 rad/s (22036 x *g*). The resulting clarified lysate was applied to NiNTA resin. The NiNTA column was then washed with 3 column volumes (CVs) lysis buffer and HisSUMO-YcnIΔC was eluted in 150 mmol/L NaCl, 20 mmol/L HEPES, 250 mmol/L imidazole, pH 7.5. To cleave the HisSUMO tag, the eluate was incubated with His-Ulp1 protease in a lysis buffer overnight with nutation at 4 °C or dialyzed against 150 mmol/L NaCl, 20 mmol/L HEPES, pH 7.5. The untagged YcnIΔC protein was further purified by applying the cleavage products to NiNTA resin and collecting the flow-through, which was then concentrated in 10 kDa MWCO centrifugal concentrators (Pall). A final step of purification was performed using size exclusion chromatography in a buffer comprised of 150 mmol/L NaCl, 20 mmol/L HEPES pH 7.5. Peak fractions were pooled and concentrated using centrifugal concentrators. Protein concentration was measured either by absorbance at *A*_280_ using an extinction coefficient of 34950 L mol^-1^ cm^-1^ or by the Bradford assay.

### Inductively coupled plasma – mass spectrometry (ICP-MS)

Copper loading was carried out by incubating purified apo *Bs*YcnIΔC with 1 mole equivalent to 2 mole equivalents of CuSO_4_ on ice for two hours prior to desalting using a desalting column. The concentrations of the desalted proteins were measured by absorbance at 280 nm (*A*_280_). Both the copper-loaded samples and the apo protein samples were prepared for ICP-MS by diluting them to 0.1 µmol/L to 0.2 µmol/L in 0.239 mol/L (1 % v/v) nitric acid. The ICP-MS was optimized for abundance sensitivity under hot plasma conditions with a 1 mL/min glass microconcentric nebulizer. Copper quantification followed measurement of ^65^Cu in pulse counting mode over five passes and three runs with 100 ms dwell time. The instrument calibration spanned less than two orders of dynamic range with five matrix matched standards bracketing the measured copper concentrations. Calibration standards from two separate vendors were diluted from primary standards and verified against each other within the same run. Repeated analysis of unknown samples agreed within ≈ 2 % and replicate experiments within ≈ 9 %. All quoted uncertainties are one stand deviation statistical uncertainties from multiple repeat measurements, unless noted otherwise.

### Electron paramagnetic resonance (EPR) spectroscopy

Purified *Bs*YcnIΔC was incubated with 1 mole equivalent of CuSO_4_ on ice for two hours. The sample was then desalted into 150 mmol/L NaCl, 20 mmol/L HEPES pH 7.5 using a desalting column. The sample was concentrated to 150 µmol/L in a buffer composed of 150 mmol/L NaCl, 4.1 mol/L (30 % v/v) glycerol, and 20 mmol/L HEPES, pH 7.5. A 150 µmol/L sample of apo protein for EPR studies was prepared in the same buffer. Samples were placed in pre-cooled EPR tubes before being capped and frozen to 77 K in liquid nitrogen. Spectra were collected at (10.0 ± 0.1) K on a commercial spectrometer operating at 9 GHz using a liquid He flow-through cryostat for continuous wave EPR measurements. A power saturation series was collected to confirm that the experimental spectra presented here were collected under non-saturating conditions. The *g* values and hyperfine coupling constants (*A*, in MHz) reported in the manuscript were determined directly from the spectra. From the spectra, *g* values were determined at the maxima, minima, and baseline-crossing points as reported by the analysis software provided by the instrument manufacturer. Based on the manufacturer-reported field (0.08 mT) and frequency (50 kHz) accuracies and the field resolution (≈ 0.1 mT per point, calculated from sweep width/number of points *viz*. 200 mT per 2048 points), the propagated uncertainty on the *g* values is ± 0.002. Using the spectra, *A*_*ǁ*_ values were determined by measuring the difference in the field position (mT) of Cu hyperfine peak maxima at the lowest field and highest field position (that is visible) and dividing by two. The field positions of the peak maxima were determined both by a peak-picking algorithm provided in vendor instrument-supplied software and by taking the first derivative of the hyperfine peak region to determine the baseline crossing point of each peak. The two methods returned field values that agreed to within 0.1 mT for the lowest-field hyperfine peak and to within 0.3 mT for the highest-field hyperfine peak. Including the manufacturer-reported field accuracy (0.08 mT) and the field resolution of ≈ 0.1 mT per point (see above), the propagated uncertainty on *A*_*ǁ*_ is calculated to be ± 0.4 mT.

### Crystallization and structure determination of Cu-bound YcnIΔC

*Bs*YcnIΔC was concentrated to 12 mg/ml and was incubated with 0.9 mole equivalents CuSO_4_ on ice for one hour. Crystallization screens were set up at room temperature using a 1:1 protein:precipitant ratio in sitting drop trays. An initial crystallization hit was obtained in 0.1 mol/L citric acid pH 3.5 and 2.0 mol/L ammonium sulfate. Crystallization conditions were further optimized to 0.1 mol/L citric acid, 2.17 mol/L ammonium sulfate. Crystals were cryoprotected by supplementing the drop with 3.57 mol/L (20 % v/v) ethylene glycol. Data were collected at the 21-ID-D beamline at the Advanced Photon Source and processed to 2.11 Å resolution using XDS GUI in space group *P*6_3_22. The structure was solved by molecular replacement in phenix.phaser (50) using a structure of a closely related protein from *N. farcinica* (PDB ID 3ESM) as a search model (Top LLG: 57.2, Top TFZ: 8.65). AutoBuild was used to place additional residues in the initial solution, resulting in 246 residues in 2 chains. An anomalous map was generated using phenix.maps (50) to determine the placement of copper ions. The structure was further improved by iterative rounds of model building and refinement in phenix.refine (50) and Coot (51), respectively. The final model consists of 128 residues in Chain A and 126 residues in Chain B with R_work_/R_free_ = 18.04 % /20.24 %. The coordinates and structure factors have been deposited in the PDB with accession code 7MEK. All software used for structure determinations and refinements are open source.

### Crystallization and structure determination of apo YcnIΔC

Crystallization screens with the apo protein were set up at room temperature using a 1:1 protein:precipitant ratio in sitting drop trays. An initial crystallization hit was obtained in 0.1 mol/L citric acid pH 3.5 and 2 mol/L ammonium sulfate. The conditions were then optimized to 0.1 mol/L citric acid pH 3.5 and 2.394 mol/L ammonium sulfate. Crystals were cryoprotected in 3 mol/L sodium malonate prior to harvesting. Data were collected at the 21-ID-F beamline at the Advanced Photon source. The data were processed to 0.205 nm (2.05 Å) resolution in space group *P*6_3_22 using XDS GUI (52). The structure was solved by molecular replacement using the Cu bound YcnI structure as a search model (Top LLG: 662.3, Top TFZ: 38.0). The structure was further refined using phenix.refine (50) and model building was carried out in Coot (51) resulting in the final structure with R_work_/R_free_ = 19.6 %/21.9 %. We also observed positive electron density in the *F*_o_-*F*_c_ map near the N-terminus of both molecules in the asymmetric unit. We attempted to model metal ions as well as components of the crystallization condition, but all gave rise to unusually high (>1.30 nm^2^ or >130 Å^2^) B-factors and did not result in improvements to the difference map. Modeling in a malonate ion due to the use of sodium malonate as a cryoprotectant did result in improved B-factors (0.7161 nm^2^ or 71.61 Å^2^) and electron density maps. The coordinates and structure factors have been deposited in the PDB with accession code 7ME6.

## Acknowledgements

This work was supported by start-up funds and a Class of ‘68 Award from Lehigh University (O.S.F.) and by financial support of the National Institute of Standards and Technology (V.A.S.). This research used resources of the Advanced Photon Source, a U.S. Department of Energy (DOE) Office of Science User Facility operated for the DOE Office of Science by Argonne National Laboratory under Contract No. DE-AC02-06CH11357. Use of the LS-CAT Sector 21 was supported by the Michigan Economic Development Corporation and the Michigan Technology Tri-Corridor (Grant 085P1000817). We thank the staff at the LS-CAT beamline for assistance with crystallographic data collection. We also thank Damien Thévenin, K. Jebrell Glover, Jing Guo, and Aarshi Singh for helpful comments and suggestions on the manuscript.

## Tables and Figures

**Table S1.**
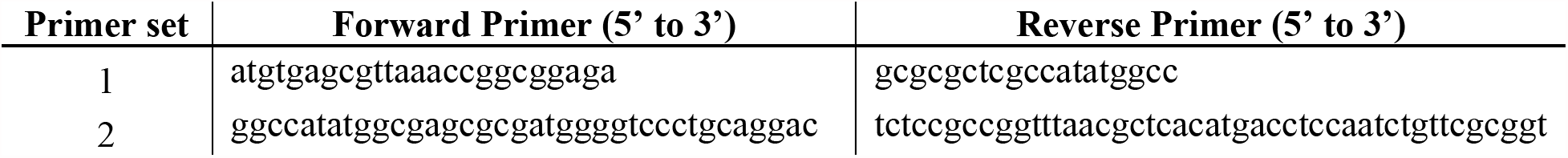
Primers used in construct design.

**Figure S1.**
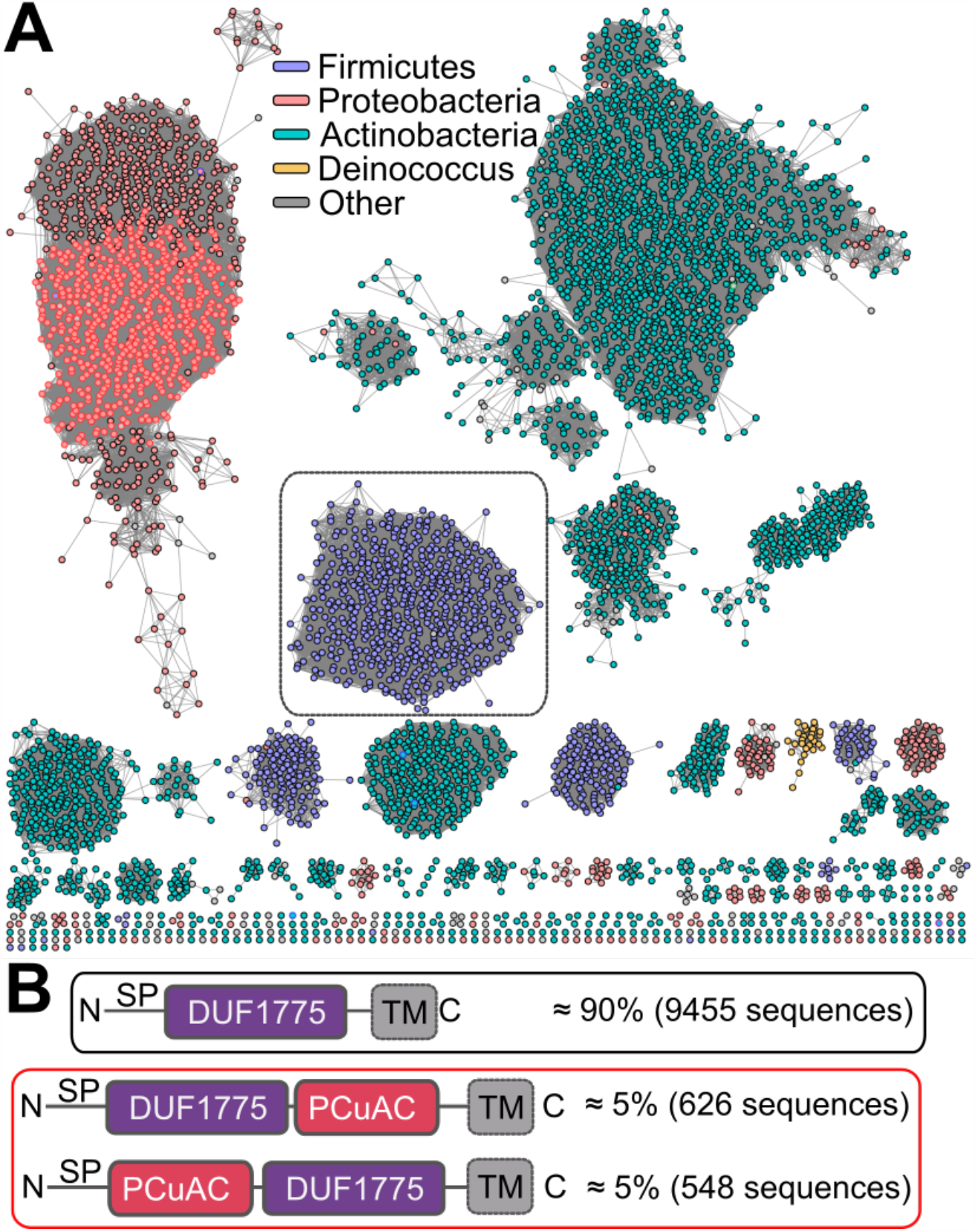
Bioinformatics analysis of all DUF1775 sequences. **A)** Sequence similarity network of 10,646 DUF1775 domain-containing protein sequences identified through JGI-IMG, colored by taxonomic information, and using an alignment score cut-off of 50. Sequences outlined in black contain only the DUF1775 domain; sequences outlined in red are fusions with PCu_A_C domains. **B)** Schematics of the most common domain organizations, with outlines using same color scheme as described in A, above. SP, signal peptide; TM, transmembrane helix.

**Figure S2.**
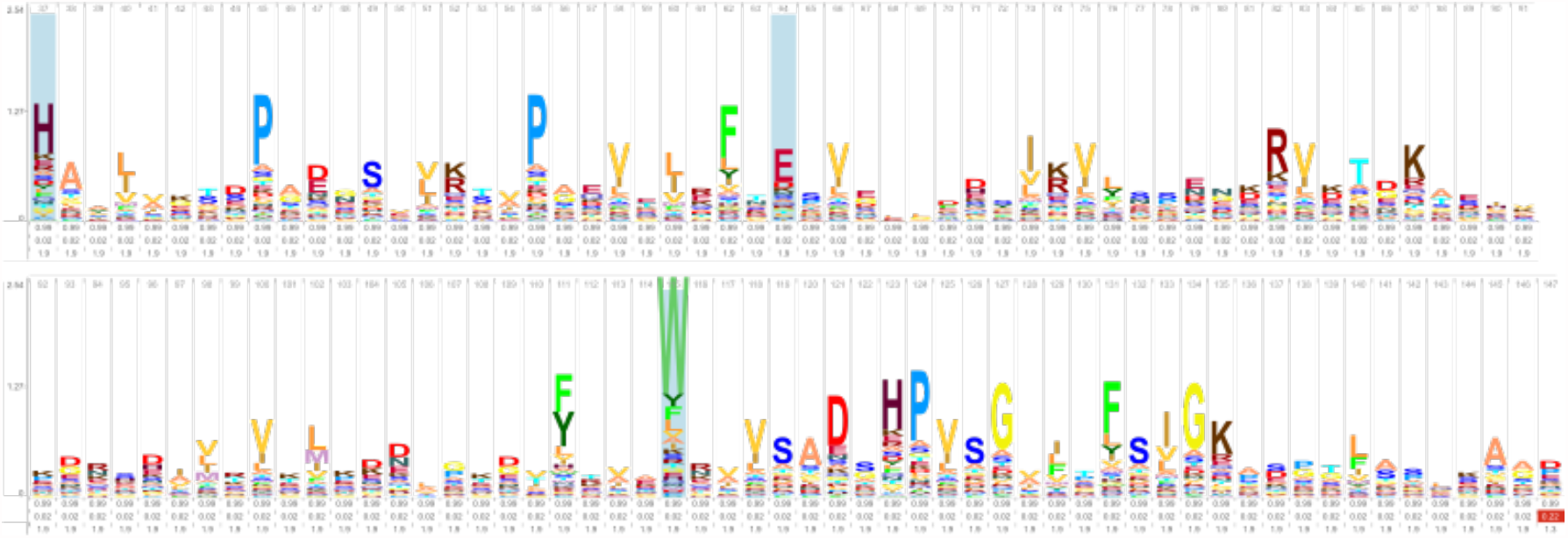
DUF1775 domain sequence logo. Sequence logo for all DUF1775 sequences used in the sequence similarity network. The positions of the three residues at the metal binding site (His, Glu, and Trp) are highlighted in blue.

